# DeeReCT-TSS: A novel meta-learning-based method annotates TSS in multiple cell types based on DNA sequences and RNA-seq data

**DOI:** 10.1101/2021.07.14.452328

**Authors:** Juexiao Zhou, Bin Zhang, Haoyang Li, Longxi Zhou, Zhongxiao Li, Yongkang Long, Wenkai Han, Mengran Wang, Huanhuan Cui, Wei Chen, Xin Gao

**Affiliations:** Computational Bioscience Research Center (CBRC), King Abdullah University of Science and Technology (KAUST), Thuwal, Saudi Arabia; Department of Biology, School of Life Sciences, Southern University of Science and Technology, Shenzhen, China; Shenzhen Key Laboratory of Gene Regulation and Systems Biology, School of Life Sciences, Southern University of Science and Technology, Shenzhen, China; Academy for Advanced Interdisciplinary Studies, Southern University of Science and Technology, Shenzhen, China

**Author notes:** co-first authors (Juexiao Zhou, Bin Zhang).

## Abstract

The accurate annotation of transcription start sites (TSSs) and their usage is critical for the mechanistic understanding of gene regulation under different biological contexts. To fulfill this, on one hand, specific high-throughput experimental technologies have been developed to capture TSSs in a genome-wide manner. On the other hand, various computational tools have also been developed for *in silico* prediction of TSSs solely based on genomic sequences. Most of these computational tools cast the problem as a binary classification task on a balanced dataset and thus result in drastic false positive predictions when applied on the genome-scale. To address these issues, we present DeeReCT-TSS, a deep-learning-based method that is capable of TSSs identification across the whole genome based on both DNA sequences and conventional RNA-seq data. We show that by effectively incorporating these two sources of information, DeeReCT-TSS significantly outperforms other solely sequence-based methods on the precise annotation of TSSs used in different cell types. Furthermore, we develop a meta-learning-based extension for simultaneous transcription start site (TSS) annotation on 10 cell types, which enables the identification of cell-type-specific TSS. Finally, we demonstrate the high precision of DeeReCT-TSS on two independent datasets from the ENCODE project by correlating our predicted TSSs with experimentally defined TSS chromatin states. Our application, pre-trained models and data are available at https://github.com/JoshuaChou2018/DeeReCT-TSS_release.

## Introduction

The transcription start site (TSS) is where the first nucleotide of a gene is transcribed (Danino, Even, Ideses, & Juven-Gershon, 2015). Promoter regions around TSS usually contain multiple *cis*-elements recognized by polymerase-recruiting transcription factors (Konoshita et al., 2004). Therefore, to precisely locate TSS and its associated promoter region is essential for understanding the *cis-trans* networks of transcriptional regulation (Triska, Ivliev, Nikolsky, & Tatarinova, 2017).

To fulfil this, specific high-throughput experimental technologies have been developed to capture TSSs in a genome-wide manner. Taking advantage of the fact that mature RNA transcripts produced by polymerase II (Pol II) have a specific cap structure at their 5’end, a so-called CAGE (Cap Analysis of Gene Expression) method has been developed to map TSS positions as well as usage in a quantitative manner. By integrating the CAGE method with massive parallel sequencing (CAGE-seq), it can capture TSSs in a genome-wide manner. In the last two decades, Functional Annotation of The Mammalian Genome (FANTOM), a worldwide collaborative project aiming at identifying all functional elements in mammalian genomes, have performed CAGE-seq in dozens of cell lines and primary tissues (Forrest et al., 2014). In total, it has collected 201,802 robust CAGE peaks for TSS annotation in the human genome. At the chromatin level, promoter regions are enriched with specific histone post-translational modifications, including H3K4me3, H3K9ac and H3K27ac. In addition, DNA around TSS is usually unbound by nucleosome, therefore manifested as a nucleosome depletion region (NDR). Together, such NDR flanked by regions enriched in H3K4me3, H3K9ac and H3K27ac modifications could be used to represent a promoter-specific chromatin state (Barth & Imhof, 2010). In the Encyclopedia of DNA Element (ENCODE) project, chromatin immunoprecipitation followed by high throughput sequencing (ChIP-seq) has been applied to characterize a series of histone modification patterns in the human genome. Using a hidden Markov model, these data enable the prediction of different chromatin states, including the promoter-specific one (Ernst & Kellis, 2012).

In addition to the experimental approach, many computational methods have been developed to predict TSSs by learning features in the flanking regions of the annotated TSSs. Several canonical elements have been identified within the promoter region around TSSs, such as the TATA-box (Osborne, Schell, Burch-Jaffe, Berget, & Berk, 1981), TFIIB recognition element (Deng & Roberts, 2005), and Initiator (Smale & Baltimore, 1989). However, studies also show that the promoter structure is highly diverse and sequence features for determining promoters are often not universal, which indicates the difficulty to globally predict promoters and TSSs in the genome. For example, it has been shown that only 17% of eukaryotic core promoters have TATA-box (Yella & Bansal, 2017). Therefore, traditional methods for promoter prediction can only detect about 50% of promoters from the genome with a high rate of false positives (Shahmuradov & Solovyev, 2015).

To address this, two major strategies have been employed to improve the performance of promoter/TSS prediction. One focuses on manual extraction of TSSs and promoters related *cis*-elements, and improves the performance by extending the predefined feature sets. For example, TSSG and TSSW (V. Solovyev & Salamov, 1997) consider the TATA-box score, *cis*-element preferences, and potential trans-factor binding sites (Wingender, 1994). PromH extends the feature set of TSSW by taking into account conserved features of major promoter functional components, including transcription start points, TATA-boxes and regulatory motifs, in pairs of orthologous genes, to further improve the promoter/TSS prediction accuracy (V. V. Solovyev & Shahmuradov, 2003). The second strategy applies machine learning models to automatically extract features to predict promoter/TSS. Promoter2.0 is the seminal work that uses neuron networks in the promoter identification (Knudsen, 1999) and DragonGSF further uses the information of GC contents and CpG islands together with neural networks to identify promoter regions (Bajic & Seah, 2003). Recently, deep-learning has been applied to the task due to the availability of massive datasets and its power of automatically extracting features. These convolutional neural network (CNN) based methods, including PromCNN (R. K. Umarov & Solovyev, 2017), TSSPlant (Shahmuradov, Umarov, & Solovyev, 2017), PromID (R. Umarov, Kuwahara, Li, Gao, & Solovyev, 2019), TransPrise (Pachganov et al., 2019) and iPSW(PseDNC-DL) (Tayara, Tahir, & Chong, 2020) significantly outperform previous traditional manual feature set-based tools on their target problems. As one of the latest developments, iPSW(PseDNC-DL) (Tayara et al., 2020) mainly focuses on the prokaryotic promoter prediction and reports good performance. To achieve the precise prediction of TSS positions, TransPrise (Pachganov et al., 2019) reports the ability to precisely predict the TSS positions by taking a two-step strategy (firstly determine whether a TSS is present or not in a long fragment, then map the position of TSS in the fragment). Importantly, the majority of machine learning-based methods for TSS prediction can only solve the binary classification task on balanced data and cannot be applied for genome scanning due to the extreme data imbalance (i.e., the ratio between non-TSS and TSS is about 15000 as 3 billion bases human genome only contain ~200,000 TSSs). So far, only PromID is able to scan the TSS in the small regions around known TSS (−5k~5k) and claims the ability to scan promoter/TSS on a genome-scale.

More importantly, despite the fact that the DNA sequence of a promoter is identical across different types of cells, it might be activated in only certain cell types, which shapes a cell-type-specific TSS landscape. However, all of the previous computational methods only consider the DNA sequence information and therefore are apparently unable to determine whether a TSS is active or inactive in a given cell type. Although experimental CAGE-seq or chromatin state-based methods (see above) are able to identify active TSSs, they are rather laborious and costly to perform on a grand scale. As a result, other than FANTOM, ENCODE and a couple of epigenome roadmap projects, the CAGE-seq or histone-modification data has only been sparsely collected. In contrast, conventional RNA-seq has been routinely used to characterize transcriptional profile across tremendous amounts of primary tissues and cell lines. Thus, for computational methods to predict active TSSs in a biological sample, it would be convenient to integrate the conventional RNA-seq data. On one hand, RNA-seq data would provide a distinct coverage pattern at a TSS flanking region. Theoretically, across a TSS, there should manifest a sharp increase of RNA-seq coverage. On the other hand, since only expressed regions need to be scanned, by integrating RNA-seq data, the total number of sites for genome-wide TSS prediction could be dramatically reduced.

Here, we introduce DeeReCT-TSS, a novel deep-learning-based method for the accurate prediction of TSSs by incorporating both DNA sequences and RNA-seq coverage information. For any sample with conventional RNA-seq data, our method could predict active TSSs in a genome-wide manner. Furthermore, by extending our method through meta-learning, we use DeeReCT-TSS for simultaneous TSS annotations on 10 cell types and are able to identify cell-type-specific TSSs. After extracting sequence features from the model, putative TFs regulating these cell-type-specific TSSs could be identified. Finally, we validate the generalization power of the model by showing that the predictions from two independent ENCODE datasets are highly consistent with the TSSs identified based on chromatin states.

## Results

### A deep-learning-based model for TSS prediction using both DNA sequence and RNA-seq coverage information

For TSS prediction, we built a deep-learning model by taking both DNA sequences and RNA-seq coverage information as inputs. In brief, for each site/nucleotide (nt), the DNA sequence and the RNA-seq coverage in the 1000bp flanking window were converted into a 1000×4 (one-hot encoding) and 1000×1 vector, respectively. Both inputs went through two independent convolutional neuron network (CNN) models that have a convolution layer with 64 filters to extract motif features and the trend of coverage changes, respectively. Following the convolution layer, a rectified linear unit (ReLU) was applied as the activation function followed by the max-pooling layer. Two feature vectors were then concatenated together and fed into the fully-connected (FC) layer. The last softmax layer gave the prediction value between 0 and 1. To improve the generalization capability weight decay and dropout were applied (Methods and Fig. 1a).

**Figure 1:**
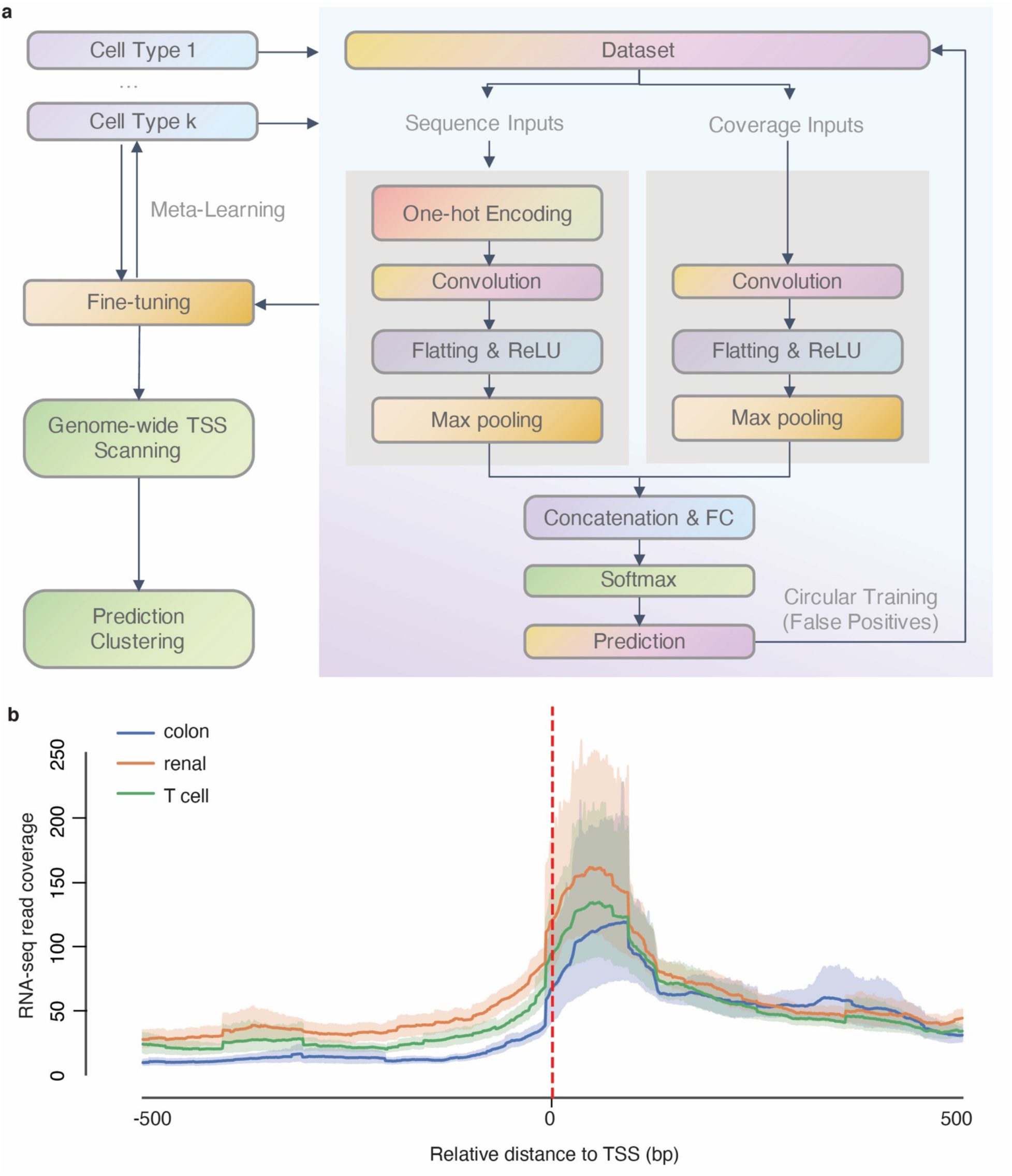
A deep-learning-based model for TSS prediction using both DNA sequence and RNA-seq coverage information. a) The schematic of the workflow and architecture of DeeReCT-TSS. b) RNA-seq coverage is dramatically increased across the TSS. The dashed line indicating the position of TSS, the solid curve is the mean of RNA-seq converge across all true TSSs and the shade is the 95% confidential interval of the coverage. Colon: The colon carcinoma cell line; Renal: The renal carcinoma cell line; T cell: The adult T-cell leukemia cell line.

### Incorporating RNA-seq coverage information improves TSS prediction

To train our model, we downloaded the full set of annotated TSSs (n = 201,802) from FANTOM (https://fantom.gsc.riken.jp/5/), and then calculated their usage in three cell lines, including a colon carcinoma (COLO-320), a renal carcinoma (OS-RC-2) and a T-cell leukemia cell lines (ATN-1), based on the respective CAGE-seq data (Methods). The active TSSs (CAGE-seq > 5 and RNAseq RPM > 0.5 or CAGE-seq > 15) from protein-coding genes were used as the positive dataset, with their corresponding DNA sequences and RNA-seq coverages as inputs. An average of ~20000 active TSSs from on average 11000 protein-coding genes was obtained in each cell line (Supp Table S1). By integrating RNA-seq data into the model, we could capture the change of the sequencing coverage around the TSS (Fig. 1b). To compare our integrative model with that using only DNA sequences or RNA-seq coverage information alone, we did the ablation study and trained three models (the integrated model, the sequence only model and the coverage only model) independently on the same training dataset and tested the dataset with identical initial weights, biases, and number of epochs (Methods). Apparently, the integrative model achieves the best performance (94%, 90%, and 92% F1-score in the colon carcinoma, the renal carcinoma and the T-cell leukemia cell lines, respectively), which is on average 3% and 13% better than the sequence only model and coverage only model, respectively (Supp Table S2).

### DeeReCT-TSS outperforms three state-of-the-art methods in the binary classification of promoter/TSS

To further evaluate our method, we selected three state-of-the-art (SOTA) methods on the promoter/TSS identification for the comparison, including PromID (R. Umarov, Kuwahara, Li, Gao, & Solovyev, 2019), iPSW(PseDNC-DL) (Tayara, Tahir, & Chong, 2020), and TransPrise (Pachganov et al., 2019). We trained each model with a very large number of epochs until (almost) convergence on the validation dataset (Methods). As shown in Table 1, our method outperforms all these three methods in terms of accuracy, TPR, FDR and F1 score, and achieves, on average, 3%, 6% and 12% improvement on F1 score compared to PromID, TransPrise and iPSW(PseDNC-DL) respectively. Since PromID is the one with the closest performance to our method (Table 1) and it is also the only published method that can control the false positives in a genome-wide scanning task, we only kept PromID for the following comparison in the genome-scale scanning task.

**Table 1:**
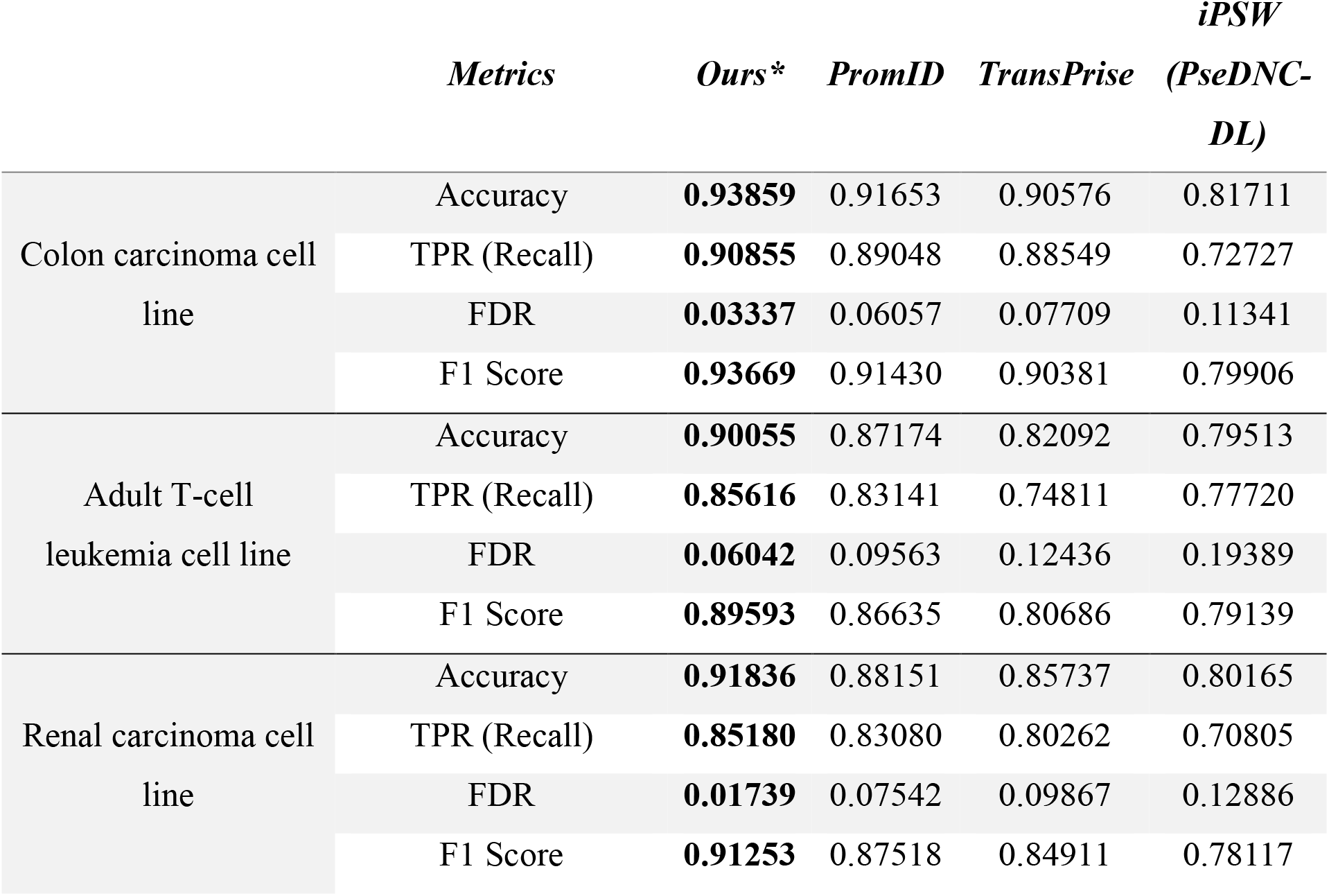
Comparison of the performance between DeeReCT-TSS and other three recently published methods in binary classification of TSSs on three datasets.

**Table 2:** Performance of meta-learning model and fine-tuned model on 10 cell lines.

|  | *Metrics* | *TPR (Recall)* | *FDR* | *F1-Score* |
| --- | --- | --- | --- | --- |
| Meta-Model | NK T-cell leukemia cell line | 0.72730 | 0.03287 | 0.83024 |
|  | Colon carcinoma cell line | 0.86235 | 0.02485 | 0.91529 |
|  | Gastrointestinal carcinoma cell line | 0.75460 | 0.03171 | 0.84819 |
|  | Acute lymphoblastic leukemia cell line | 0.71928 | 0.02943 | 0.82624 |
|  | Lung carcinoma cell line | 0.74477 | 0.03022 | 0.84251 |
|  | Myeloma cell line | 0.75222 | 0.02814 | 0.84805 |
|  | Neuroblastoma cell line | 0.79193 | 0.02486 | 0.87403 |
|  | Renal carcinoma cell line | 0.77725 | 0.02699 | 0.86418 |
|  | Adult T-cell leukemia cell line | 0.74834 | 0.03211 | 0.84407 |
|  | Testicular germ embryonal cell line | 0.72749 | 0.03654 | 0.82901 |
| Fine-tuned Model | NK T-cell leukemia cell line | 0.82357 | 0.05374 | 0.89206 |
|  | Colon carcinoma cell line | 0.92227 | 0.04101 | 0.94027 |
|  | Gastrointestinal carcinoma cell line | 0.86808 | 0.05310 | 0.90578 |
|  | Acute lymphoblastic leukemia cell line | 0.85455 | 0.06530 | 0.89282 |
|  | Lung carcinoma cell line | 0.86677 | 0.05926 | 0.90224 |
|  | Myeloma cell line | 0.87594 | 0.05743 | 0.90803 |
|  | Neuroblastoma cell line | 0.87728 | 0.04078 | 0.91641 |
|  | Renal carcinoma cell line | 0.87969 | 0.04738 | 0.91470 |
|  | Adult T-cell leukemia cell line | 0.86289 | 0.05618 | 0.90154 |
|  | Testicular germ embryonal cell line | 0.88168 | 0.07335 | 0.90360 |

### DeeReCT-TSS enables genome-wide promoter/TSS identification by scanning the transcribed genome

Using the RNA-seq coverage information, we could reduce the number of sites for the genome-wide TSS scanning from 3 billion to ~800 million (i.e., any site covered by more than one RNA-seq read and its flanking 1000bp), although even this reduced number of testing sites is well beyond the capacity of the previous TSS prediction methods to achieve a reasonable number of false positives.

Since positive and negative datasets are highly unbalanced during the genome scanning, in which less than 0.01% of scanning sites are true TSSs and the rest are all negatives, we introduced an iterative negative data enhancement strategy into our model. In brief, we repeatedly trained the binary classification model by randomly substituting negative data in the training dataset with false positives predicted by the model trained in the previous round (Methods and Fig. 1). By doing so, the model was expected to see more and more difficult negative samples (i.e., the ones that are more and more similar to the positive ones), therefore its power in distinguishing the true positives from the negatives would be boosted.

In the genome-wide scanning, we predicted TSS scores for 837,507,571, 892,712,017 760,462,470 sites covering 15517, 15982, 15736 protein-coding genes in the colon carcinoma, the renal carcinoma and the T-cell leukemia cell lines, respectively. In all of these three datasets, compared to PromID, our method could achieve a higher recall with a much lower false-positive rate (FPR) (Fig. 2a). However, even though the FPR of both methods are relatively low (with a 90% recall rate, FPR of DeeReCT-TSS: 8-10 FP per kb, PromID: 20-30 FP per kb), due to a total number of ~800 million query positions, millions of sites predicted as TSSs are still false positives. To address this, we further grouped the sites into clusters based on their prediction scores and used the score distribution in each cluster to evaluate whether it contains a true TSS or not (Methods and Fig. 1). As we expected that sites in the vicinity of the true TSS should get high scores, the clusters harboring the true TSS should contain a very dense set of highly-scored-site. In contrast, false positives could be derived from sparsely distributed high-score-sites. By taking this into account, we could transform predictions in ~800 million sites to less than one million clusters.

**Figure 2:**
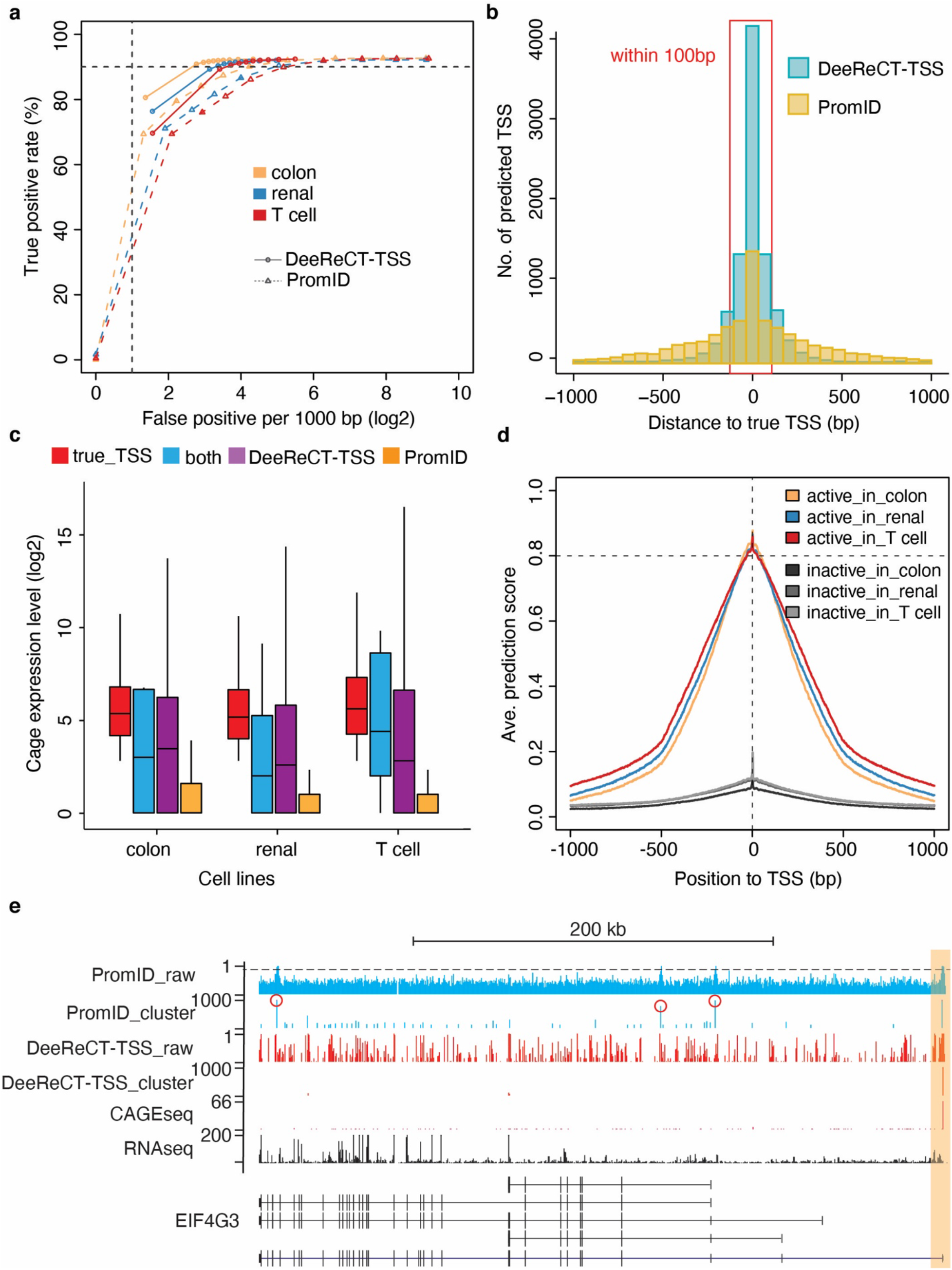
DeeReCT-TSS outperforms a recently published method in TSS identification. a) The performance of DeeReCT-TSS and PromID on datasets from three cell lines. b) Histogram of the distance between predicted TSSs and true TSSs in the colon carcinoma cell line. c) Boxplot showing the CAGE expression of multiple groups of TSS, including: DeeReCT-TSS: TSSs predicted only by DeeReCT-TSS; PromID: TSSs predicted only by PromID; both: TSSs predicted by both DeeReCT-TSS and PromID. true_TSS: ground truth in each cell line, which is the actively expressed annotated TSSs. d) Metagene analysis of active and inactive TSSs with the prediction score from DeeReCT-TSS in three cell lines. e) Genome browser view of CAGE-seq, RNA-seq, raw prediction scores outputted from DeeReCT-TSS (DeeReCT-TSS_raw) and PromID (PromID_raw), and scores after clustering (DeeReCT-TSS_cluster and PromID_cluster) in gene EIF4G3. TSSs falsely predicted by PromID were marked with red circles.

Based on the total number of true and false predicted TSSs, for each site, we could estimate an empirical probability of this site to be a true/false TSS. For instance, for any site with a prediction score of 0.1, the probability to be a true TSS is on average 2.9% among all sites with a score larger than 0.1 from the three cell lines. Then for each cluster, using a null hypothesis that there is no true TSS within this cluster, we were able to calculate this probability, which is the p-value to reject the null hypothesis and accept the alternative hypothesis that this cluster contains true TSSs (Methods). In this way, we could reduce FDR dramatically, achieving ~77% recall with ~25% FDR in the colon carcinoma cell line (Supp Fig. S1a and S1b). Notably, among the false positive predictions, 33% of them are still annotated in FANTOM and supported by CAGE-seq, but their expression levels do not pass our expression threshold. 14% are annotated but not supported by CAGE-seq and 13% of them are not annotated in FANTOM but supported by CAGE-seq. Only ~12% of the total predicted TSSs are unannotated and not supported by CAGE-seq (Supp Fig. S1a). For the false negatives, by comparing the TSS prediction in binary classification and genome scanning, we found that the sites still predicted as true TSSs in genome scanning had a higher binary classification scores than those were not identified in the genome scanning, suggesting TSSs with moderate scores in binary classification could be dropped during circular training (Supp. Fig. S1E). Moreover, the TSSs with higher expression level were more likely to be successfully predicted in genome scanning than those with lower expression (Supp. Fig. S1F).

In comparison, after the clustering procedure, to get a 70% recall, FDR of the PromID method remains as high as 50% (Supp. Fig. S1a). Moreover, the central positions of the clusters identified by our method are very close to true TSSs, which is much more accurate than PromID (Fig. 2b and Supp Fig. S1c and S1d). Finally, we could output the predicted TSSs in an extended bedgraph format (https://genome.ucsc.edu/goldenPath/help/bedgraph.html) that could be conveniently used for visualization in the genome browser as shown in Supp Table S3.

### DeeReCT-TSS could discriminate active from inactive TSSs

TSS might be only used in certain cell types and computational methods that are solely based on DNA sequences are unlikely to determine such cell-type-specific TSS usage. In comparison, by incorporating RNA-seq information, our method should be able to address this issue. To assess the performance on this perspective, we split the predicted TSSs into three groups: (a) the ones only predicted by our method, (b) the ones only predicted by PromID, and (c) the ones predicted by both methods, and compared them with true TSSs in each cell line. By analyzing the expression level of these TSSs in the respective cells, we found that the TSSs in group a and c were expressed based on CAGE-seq and the expressions were only slightly lower than that of the true TSSs, whereas the TSSs only predicted by PromID were almost not expressed at all (Fig. 2c). To further demonstrate that our method could distinguish active TSSs from inactive ones, we classified all TSSs as active or inactive based on their expression level in each cell line. We then compared the predicted score between these two groups of TSSs. As shown in Fig. 2d, only active TSSs manifested the positive signals (Fig. 2d). As one example, 24 TSSs were annotated for gene EIF4G3, while only the proximal TSS was used in the colon carcinoma cell line (represented by CAGE-seq), which could be distinguished by our method (Fig. 2e). In comparison, PromID predicted three more false TSSs, one is inactive annotated TSS and another two are even not annotated.

### A meta learning-based extension for predicting cell-type-specific TSSs and associated TFs

Since DeeReCT-TSS was able to identify active TSSs in each cell type, we would like to further explore the commonality among different cell types and, at the same time, build a model that can be easily generalized to handle the difference among them. For this purpose, the SOTA meta-learning strategy, which takes advantage of an abstract model trained with data from multiple tasks and is capable of good adaptation to a specific task with a mini fine-tuning session, exactly meets our needs. Here, the more specific aim was to train a generalized model across multiple cell types, which is also capable of being fine-tuned for any specific cell type. To check the feasibility of integrating the meta-learning strategy in DeeReCT-TSS, we extended our analysis to 10 cell lines and a total of 53,969 TSSs that are active in at least one cell type were used as the positive dataset (Material and Methods). The number of TSSs that are active across different numbers of the cell lines shows a typical bimodal distribution, where the two peaks represent those only expressed in one cell line and those expressed in all 10 cell types, respectively (Fig. 3a). Next, to obtain an abstract model with the maximum interpretability across all 10 cell types, we trained the meta-learning model with partial training data and then fine-tuned it to each cell type (Method and Fig. 1). Based on the F1 score, we found that the performance of the model using 20% of TSSs from each cell type for meta-training was saturated, while using more than 20% of TSSs had no further effect to improve the model. As shown in Table 3 and Supp Table S2, on the binary classification task, the meta-model trained with 10 cell types decreased the FDR by 1% on average compared to the model trained solely on the colon carcinoma, the renal carcinoma and the T-cell leukemia cell lines, though with on average decreased F1 score by 4% as a trade-off for the generalized model. Based on the meta-model, the fine-tuned model shows better performance than both meta-model and the one trained solely in each cell line with on average 5% and 1% improvement on F1 score respectively.

**Figure 3:**
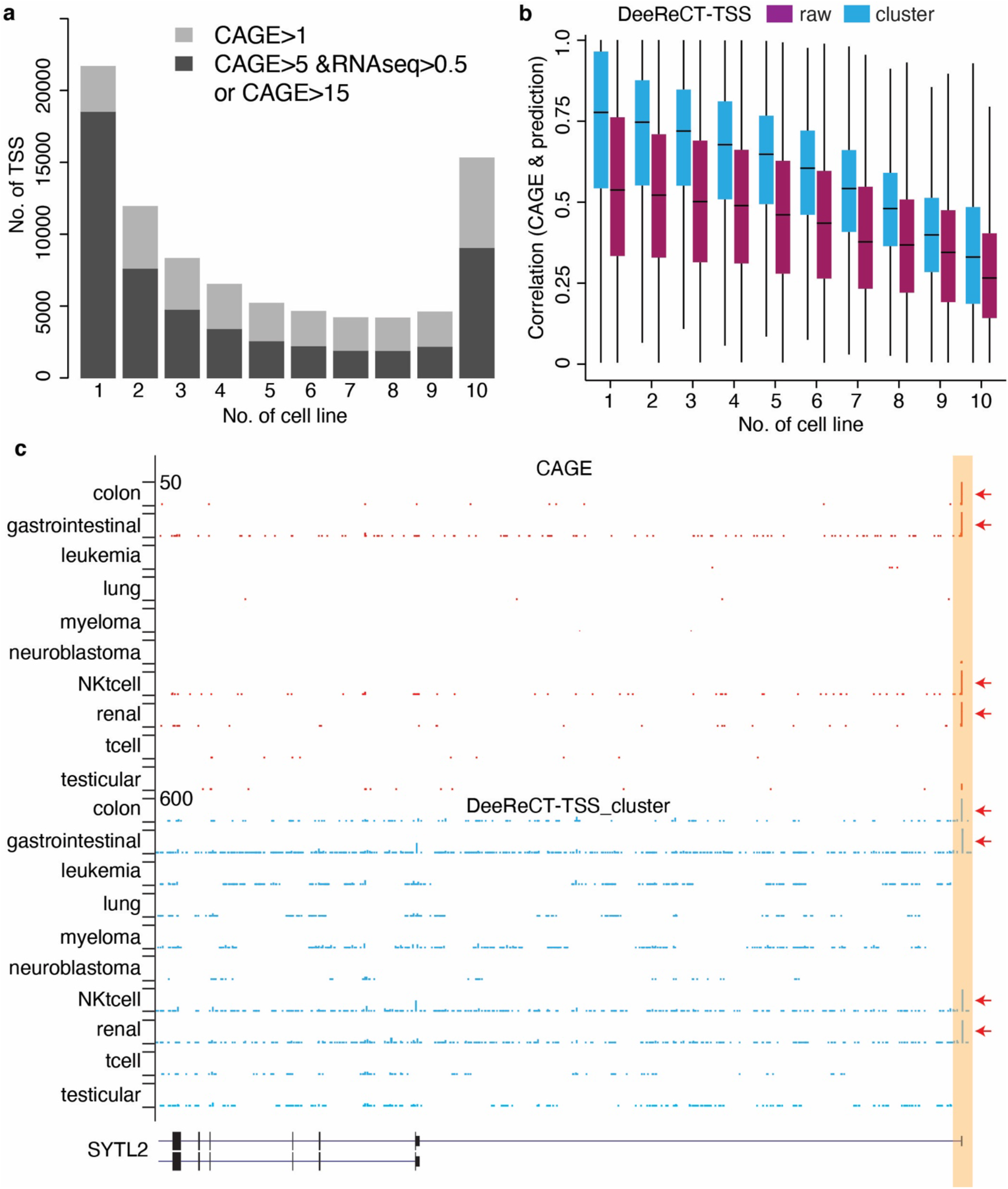
DeeReCT-TSS is capable of identifying cell-type-specific TSSs. a) Numbers of active TSS in the corresponding number of cell lines. The x-axis is the number of cell lines, and the y-axis is the number of TSS. b) Boxplot illustrating correlations between prediction scores of TSSs from DeeReCT and CAGE expressions across 10 cell lines. Each box indicating a group of TSSs expressed in corresponding cell line numbers. The raw is the raw prediction scores outputted from the model, and the cluster score is the score after clustering of the raw score. c) UCSC genome browser visualization shows an example of TSS (highlight in yellow) from gene SYTL2 that is active in the colon carcinoma, the gastrointestinal carcinoma, the NK T-cell leukemia and the renal carcinoma cell lines (represented by CAGE-seq), were correctly predicted by DeeReCT-TSS. The red arrow indicates the cell lines, where the TSS is active or predicted. The colon, gastrointestinal, leukemia, lung, myeloma, neuroblastoma, NKtcell, renal, tcell, testicular at the left side of c) are corresponding to the colon carcinoma cell line (COLO-320), the gastrointestinal carcinoma cell line (ECC12), the acute lymphoblastic leukemia cell line (HPB-ALL), the small cell lung carcinoma (NCI-H82), the myeloma (PCM6), the neuroblastoma cell line (CHP-134), the NK T-cell leukemia cell line (KHYG-1), the renal carcinoma cell line (OS-RC-2), the adult T-cell leukemia cell line (ATN-1), and the testicular germ cell embryonal carcinoma cell line (NEC15).

We further applied the fine-tuned model on the genome scanning task in each cell line, and eventually, we identified TSSs with a TPR around 73% and an FDR around 28%. Among these false predicted TSSs, half of them is annotated but the expression level does not pass the threshold, while only 10% of the total predicted TSSs is neither annotated nor supported by CAGE-seq (Supp Fig. S2a). We also compared the results from the fine-tuned model to that from the model solely trained on the colon carcinoma cell line, which shows a better performance in terms of both recall (79.5% and 77.7%, respectively) and FDR (24.4% and 25.5%, respectively). To further evaluate the performance of our model to accurately predict the TSSs with specific expression patterns across 10 cell types, we correlated the prediction scores with the CAGE-seq based expression level across 10 cell lines for each TSS and obtained a strong positive correlation (median of Pearson correlation is 0.58). Notably, the correlation between prediction and CAGE-seq expression is the lowest (median of Pearson correlation is 0.24) for those TSSs that are active in all 10 cell lines, which is likely due to the fact that the prediction score can only determine if a TSS is active or not, but it is not a metric to measure the TSS expression level. In contrast, for those cell-type-specific TSSs expressed in only one cell line, the median correlation coefficient is as high as 0.75 (Fig. 3b). Fig. 3c shows an example in gene SYTL2, which is active in the colon carcinoma, the gastrointestinal carcinoma, the NK T-cell leukemia and the renal carcinoma cell lines. Such usage pattern was successfully predicted by our method. Overall, these results suggest that our model could be generalized to identify active TSSs in different cell types.

One major advantage of our meta-learning model is that its abstract model can capture common features of TSSs among all cell lines, while the fine-tuned model to each cell line has a preference to identify the active TSSs in the corresponding cell line by enhancing the usage of the cell type specific features. To this end, we visualized the gain of motif information (GMI) from the convolutional layers by comparing the difference between paired filters of the meta-model and the fine-tuned model of each cell type and matched those GMI to the motifs of transcription factor (TF) binding sites (Methods). In total, 42 putative TFs with concordant expression patterns (Pearson correlation coefficient ≥ 0.5) were identified across 10 cell lines (Fig. 4a and 4b). These TFs that are specifically expressed in some cell types, potentially regulate the corresponding cell-type-specific TSSs. For example, BHLHA15/MIST1 identified in myeloma has been reported as a plasmacytic differentiation marker and potentially controlled transcriptional network with stage-specific overexpression during plasma cells differentiation (Kassambara et al., 2021) (Yeung et al., 2012). ETV5 identified in colon carcinoma cell lines has been reported to be abnormally upregulated in the colorectal cancer and positively correlated with the tumor size, lymphatic metastasis, tumor node metastasis and worse survival (Cheng et al., 2019). Examples of the TF motifs and the matched *cis*-elements extracted from deep learning model in different cell lines are shown in Fig. 4c.

**Figure 4:**
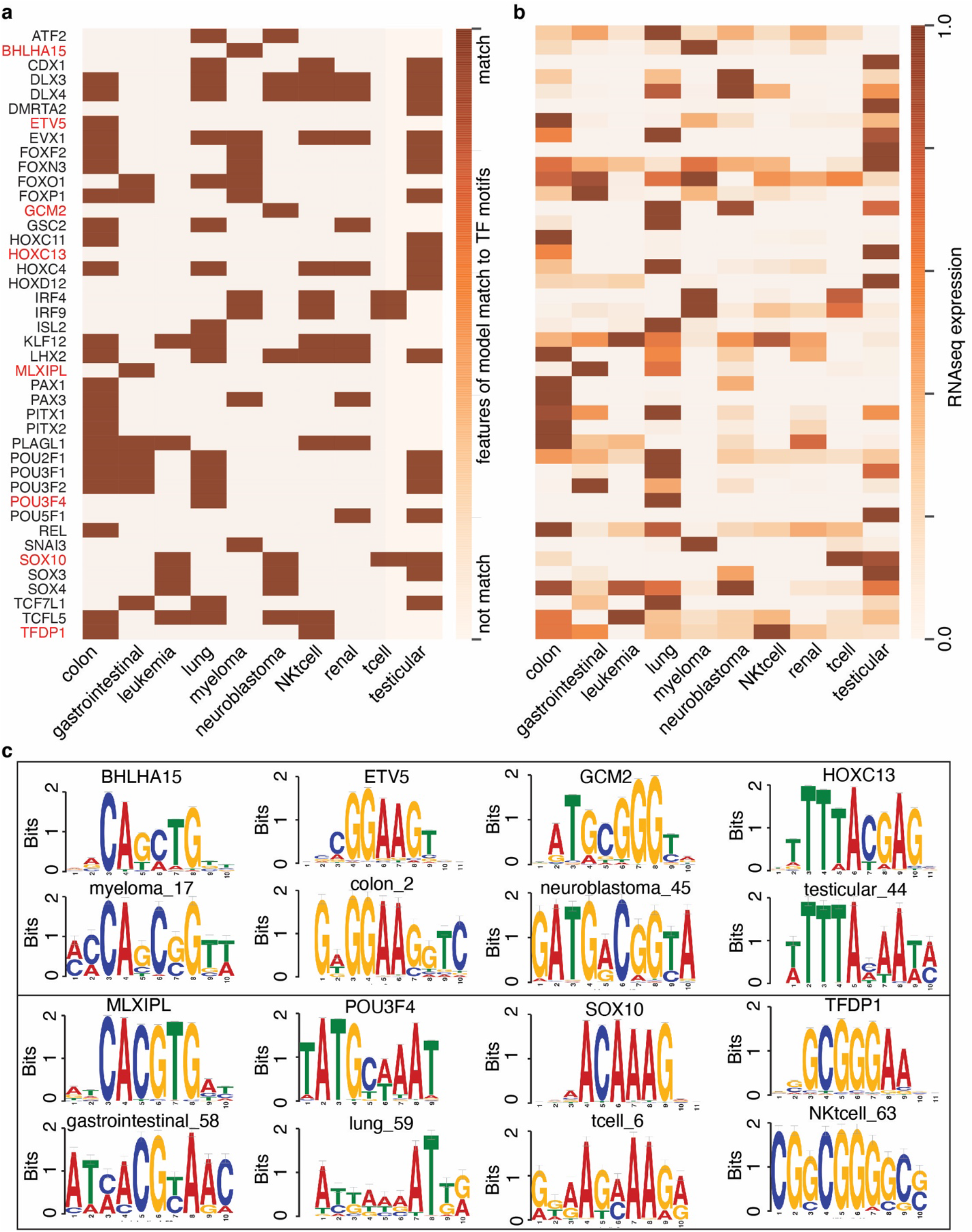
Identification of putative *cis*-elements around TSS and the corresponding transcription factors (TFs) across 10 cell lines. a) Heatmap showing that the identified TFs matched with sequence elements extracted from the deep learning model in each cell line. b) Heatmap showing the normalized expression level of the TFs across 10 cell lines. c) Examples showing the binding motifs of the identified TFs and sequence elements extracted from the deep learning model from different cell lines (marked in red in Fig. 4a).

### The trained meta-learning model could be applied to other independent datasets

As the information for the active TSSs could also be encoded at the chromatin level, to further evaluate the generalization ability on TSS identification of our meta-learning extended model to unseen datasets, we applied it to two cell lines HepG2 and K562, for which ENCODE project collected both RNA-seq and different histone modification data. In total, we predicted 14569 and 13465 putative TSSs in HepG2 and K562, respectively. Next, we associated the prediction with chromatin states of these two cell lines, which were identified by using a series of histone modification and transcription factor ChIP-seq data in the ENCODE project, including Tss, Tssf (TSS flanking region), promoterP (inactive promoter), and so on. The results showed that our predictions were highly consistent with Tss chromatin state in these two cell lines. Importantly, the overlap between the predicted TSSs in HepG2 and the Tss chromatin state from K562, as well as the overlap between the predicted TSSs in K562 with the Tss chromatin state from HepG2, were much lower than that within the same cell type, suggesting the high cell-type specificity of our prediction (Fig. 5a).

**Figure 5:**
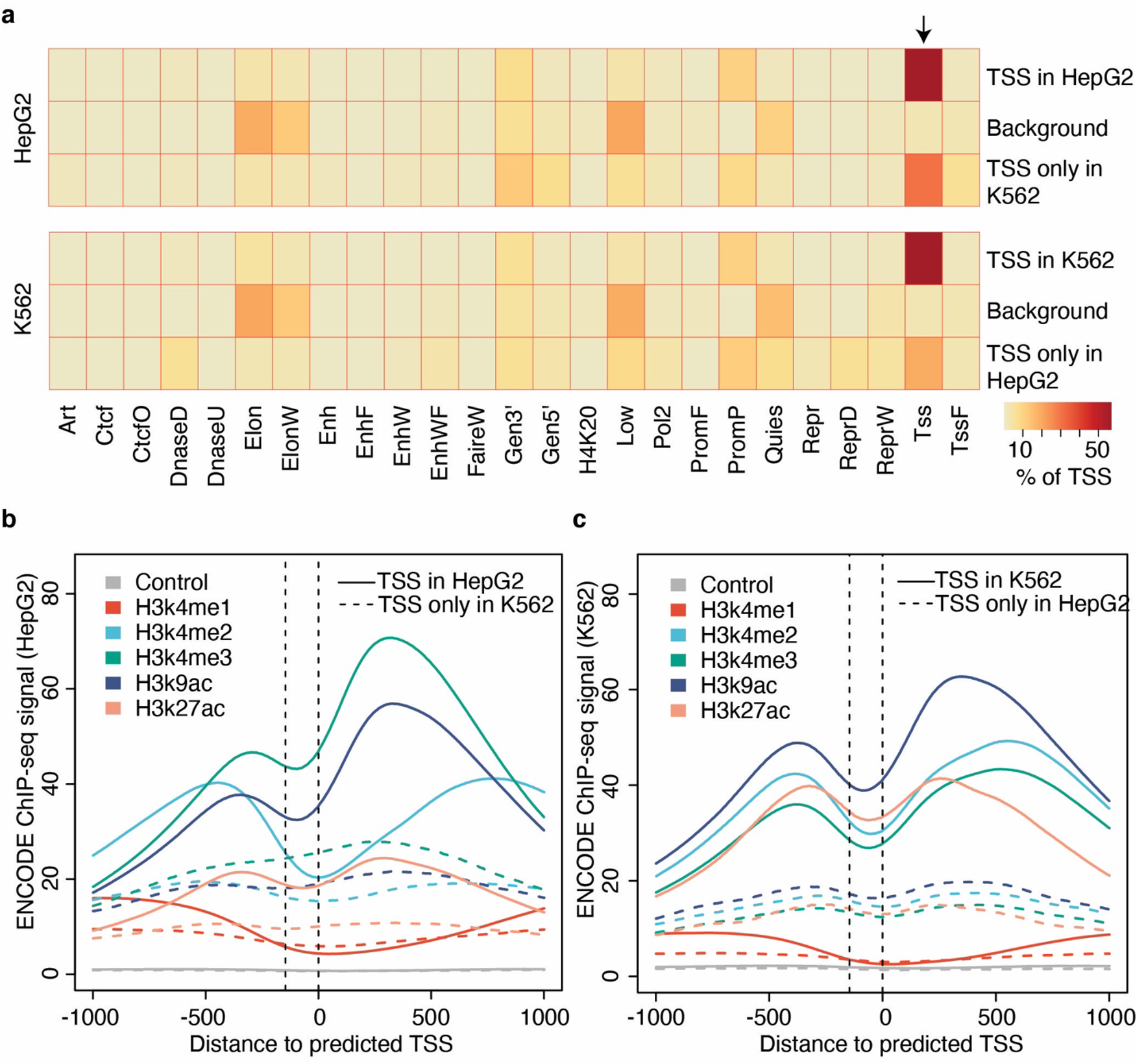
TSSs predicted in K562 and HepG2 by DeeReCT-TSS could be validated by chromatin states identified from epigenetic information. a) Heatmap showing the proportion of predicted TSSs located in each chromatin state in K562 and HepG2. The background was generated by randomly selecting regions, which were covered by RNA-seq data. b) and c) Meta-gene analysis shows that the predicted TSSs could be located within the nucleosome depletion region (NDR) in the promoter region in HepG2 (b) and K562 (c), respectively. Putative NDR within the promoter was indicated by two vertical dashed lines.

As regions of the ENCODE chromatin state are still very wide (~500bp), to inspect our predicted TSS position more precisely, we performed the meta-gene analysis of the predicted TSSs with H3K4m1, H3K4m2, H3K4m3, H3K9ac, H3K27ac ChIP-seq signals in single base-pair resolution. Surprisingly, our predicted TSSs are exactly located within the nucleosome depletion regions (NDR) flanked by the modified histones, where the transcription was likely initiated. Again, this pattern could only be observed by overlapping results from the same cell line (Fig. 5b and 5c), demonstrating again a high accuracy and specificity of our method. Taken together, all these data suggest that DeeReCT-TSS is a novel computational method that is capable of accurately predicting active TSSs in a genome-wide manner. It could be widely used to study TSS usage across multiple cell types or tissues, as well as between different pathophysiology conditions as long as conventional RNA-seq data is available.

## Discussion

Computational methods for TSS prediction have been used to annotate TSSs and to study regulatory mechanisms via *cis*-elements residing around the TSS. One shortage limiting their practical applications is that the majority of them can only do binary classification between sequences with TSS and the ones without TSS, on a balanced dataset. However, in the human genome, these positive and negative sequences are highly unbalanced, while the number of negative sequences is 15000 times more than the positive ones. In DeeReCT-TSS, we addressed this challenge by several means. First, we iteratively enhanced the negative dataset, which dramatically decreases the amount of potential false positives by forcing the model to distinguish between active TSSs and inactive TSSs with similar features. In addition, we incorporated RNA-seq data information, which significantly improves the accuracy of the predicted TSS position. Also, as we filtered out a huge number of query positions, this also helped to decrease the FDR when performing the genome scanning task. Moreover, we introduced a clustering-based method on the prediction score from the model, which further greatly reduce the FDR. Finally, meta-learning technology worked together with those strategies to ensure the robust and generalizable performance of DeeReCT-TSS in genome-scale TSS identification task.

In spite of the fact that the DNA sequence of a promoter is identical in different cell types and tissues, their transcription profiles are highly variable, resulting in distinct landscapes of active TSSs. In the last decades, the FANTOM project annotated more than 200000 TSSs in the human genome by combining CAGE-seq data from dozens of cell types, while only a handful of them (~10%) could be active in single cell type or tissue. Discriminating whether a TSS is active or inactive in each cell type or tissue could be equally important as predicting a novel TSS in studying gene regulation across cell types/tissues and between different physiology conditions. To our knowledge, DeeReCT-TSS is the only computational method that is capable of accurately identifying active TSSs in a cell type by incorporating RNA-seq data and meta-learning. By applying our method, we expect to identify more than 70% of the active TSSs and about 10% of our predicted TSSs might be true false positives.

Finally, as demonstrated in Fig. 5, the meta-model trained from the 10 cell lines, could be generalized to other independent samples with only conventional RNA-seq data (HepG2 and K562). Therefore, our model is capable of being applied to characterize active TSSs in many other cell types by using the corresponding RNA-seq data. Recently, a pan-cancer analysis using thousands of RNA-seq samples from The Cancer Genome Atlas (TCGA) project revealed the widespread alternative promoter regulations (Demircioğlu et al., 2019). However, it only studied promoters from annotated TSSs and quantified promoter activity using splicing junctions of the first exon. On one hand, there might be a tremendous amount of unannotated TSSs expressed in cancers given they often showed high abnormal transcriptional activities. On the other hand, the first splicing junctions could be hundreds and thousands bp away from TSSs, which limited the spatial resolution of that study in finding the change of TSS positions. We believe that DeeReCT-TSS could solve these problems and it could be used to identify TSSs that are specifically active in each cancer type by applying it on the TCGA RNA-seq datasets.

## Materials and Methods

### Dataset preparation

Our model has two inputs, the DNA sequence and the read coverage from RNA-seq data. DNA sequences were extracted from the human reference genome (hg19), and coverages were calculated based on RNA-seq bam files with bedtools (Quinlan & Hall, 2010) and samtools (Li et al., 2009) from the FANTOM project (https://fantom.gsc.riken.jp/5/). We firstly selected 3 cell lines to train our model, including the renal carcinoma cell line, the adult T-cell leukemia cell line, and the colon carcinoma cell line. To obtain the active TSSs in each cell line, we downloaded the robust CAGE peaks for human samples from FANTOM, which consists of 201,801 peaks. Next, for each cell line, we calculated the expression of each peak based on the downloaded homogeneous hCAGE (CAGE sequencing on Heliscope single molecule sequencer) data and RNA-seq data from FANTOM. The expressed TSSs were defined by requiring CAGE-seq score > 5 and RNA-seq coverage > 0.5 reads per million (RPM), or CAGE-seq score > 15. Finally, we only kept those TSSs located in regions (from upstream 5kb of the gene start to the gene end) of protein-coding genes. In total, an average of ~19874 TSSs were used for training in each cell line (Table 1).

To obtain the inputs for training, we calculated the coverage at each genomic site based on RNA-seq bam files. For sites within each active TSS peak, we extracted the DNA sequences and RNA-seq coverages from −500 bp to 501bp as the positive dataset. Then we randomly picked the same number of regions with a distance between 500 bp to 1000 bp from the nearest TSS peak. The full dataset was split into the training (90%) and the test dataset (10%), and in total 5 random splits were performed, and the average performance was reported.

The regions for genome scanning were defined by RNA-seq data. We extracted RNA-seq coverage at each genomic site and merged any sites covered by more than one RNA-seq read within 1000bp using bedtools.

Chromatin state segmentation of K562 and HepG2 cell lines by ChromHMM from the ENCODE project was downloaded from the UCSC genome browser (Zweig, Karolchik, Kuhn, Haussler, & Kent, 2008), while RNA-seq data was downloaded from ENCODE (“The ENCODE (ENCyclopedia Of DNA Elements) Project,” 2004) and aligned to the reference genome (hg38) with STAR (Dobin et al., 2013). To prepare the inputs for the TSS prediction, we merged 3 bam files from K562 and 4 bam files from HepG2 to get RNA-seq datasets with decent sequencing depth (~100 million reads).

### Deep neural networks for binary classification

We designed a deep neural network to extract information from both the DNA sequence and the RNA-seq coverage with two parallel CNN networks, which share a similar structure. Both inputs went through two independent CNN networks that have a convolution layer with 64 filters as the first layer to extract motif features and the trend of coverage changes. Following the convolution layer, a rectified linear unit (ReLU) was applied as the activation function followed by the max-pooling layer. Two feature vectors were concatenated together and fed into the fully-connected (FC) layer. The last softmax layer gave the prediction between 0 and 1. Weight decay and dropout were applied to improve the generalization capability of our method. We used Tensorflow-1.14.0 as the framework of our model and trained the model with one Quadro RTX 4000 GPU and on average 2 hours for 10000 epochs.

The DNA sequence was one-hot coded with dimension 1×1001×4, in which A was encoded as (1 0 0 0), T was encoded as (0 1 0 0), C was encoded as (0 0 1 0), and G was encoded as (0 0 0 1). RNA-seq coverage information was in the dimension of 1×1001×1 and directly used as the one of inputs. Both the DNA sequence and the RNA-seq coverage were fed into the network, resulting in the predicted value for each site in each TSS peak.

### Circular training for genome scanning

As binary classification is built on balanced datasets, whereas the data in genome scanning are highly unbalanced, we introduced the iterative negative data enhancement as a negative data argumentation method to our model to reduce the FPR as shown in Algorithm 1. For each repetition, we trained the model and selected the best one by evaluating with multiple metrics, including TRP, FDR and F1 score, then we applied the best model on the scanning task of randomly sampled 100 regions in the genome, whose center (+1) is the TSS used in the training dataset.

For each scanning task, we divided the large scanning window (−5000bp~5000bp) into 2 partitions, one is the positive region around the true TSS site (−500bp~+501bp, with TSS locate at +1) and all others outside the positive region were considered as negative regions. Any site predicted as TSS in the positive region will be considered as TP, otherwise FP. Among those FPs, half of the negative data in the training dataset was replaced by the randomly selected cases from scanned FPs and the whole process was repeated again for the next repetition.

We implemented this learning process with Tensorflow-1.14.0 as the framework and trained it with one Quadro RTX 4000 GPU. The average time for one repetition is 2 hours and the whole process with 20 repetitions takes 40 hours.

#### Algorithm 1 Circular training for reducing false positive

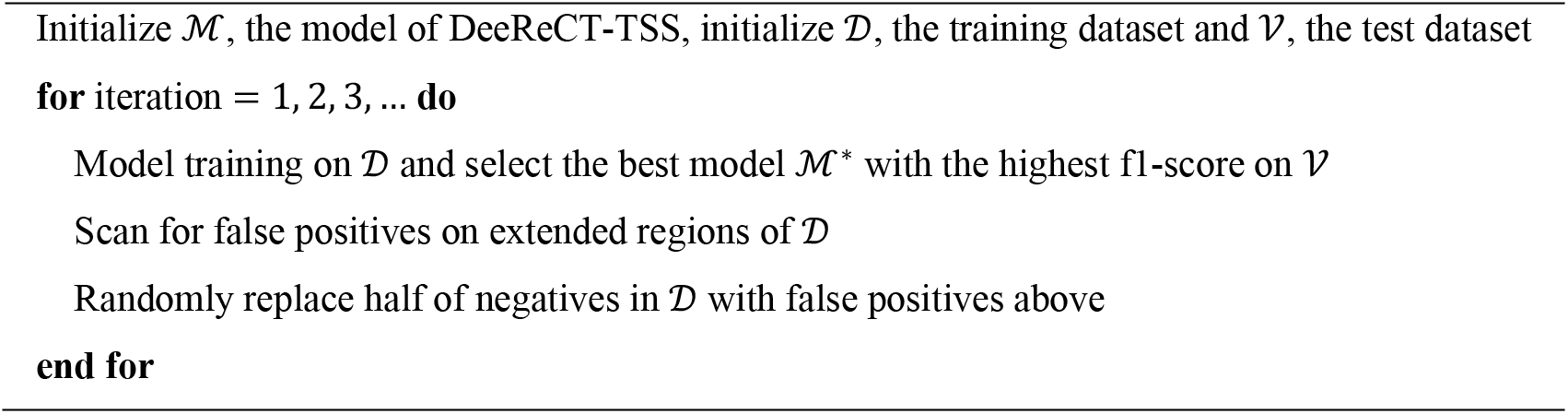

### Meta-learning across multiple cell lines

To obtain a more generalized model across different cell types, we incorporated the SOTA meta-learning algorithm and trained the meta-model from 10 cell lines, including a colon carcinoma cell line (COLO-320), a gastrointestinal carcinoma cell line (ECC12), an acute lymphoblastic leukemia cell line (HPB-ALL), a small cell lung carcinoma (NCI-H82), myeloma (PCM6), a neuroblastoma cell line (CHP-134), a NK T-cell leukemia cell line (KHYG-1), a renal carcinoma cell line (OS-RC-2), a adult T-cell leukemia cell line (ATN-1), and a testicular germ cell embryonal carcinoma cell line (NEC15). We integrated the Reptile (Nichol, Achiam, & Schulman, 2018) algorithm to achieve the meta-learning across multiple cell types as shown in Algorithm 2. Given the meta-model from 10 cell types, we further fine-tuned with 20% data of the corresponding cell type respectively and obtained the cell-type-specific model.

#### Algorithm 2 Meta-learning for DeeReCT-TSS

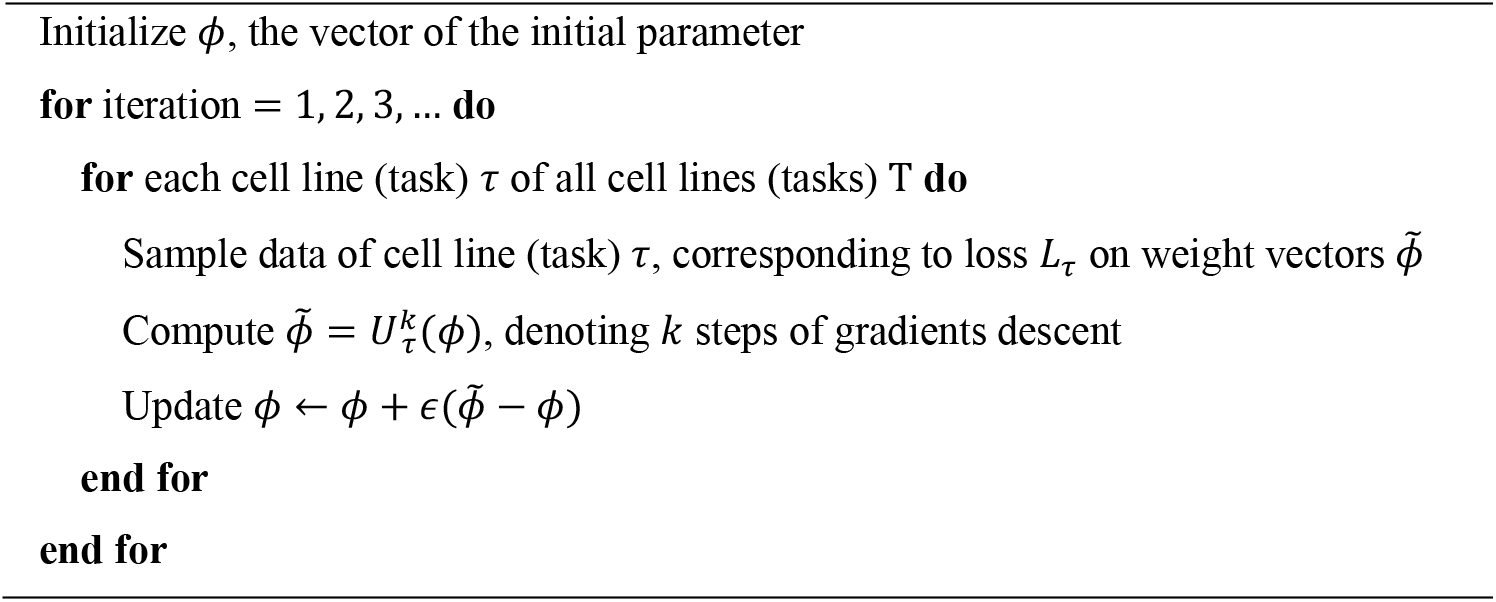

### Evaluation metrics

To evaluate our method and to objectively compare predictions by our model and other methods for TSS identification, we measured the performance using accuracy, recall, false-positive rate (FPR), false discovery rate (FDR) and F1-Score, which are defined as:

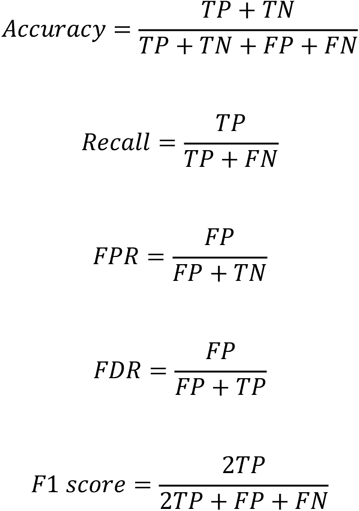

In binary classification, one region in our dataset can only be positive or negative and all metrics work without confusion. To better define the TP and FP during scanning along a certain region, we firstly searched for the strongest signal predicted by our model and counted signals with distance ≤ tolerant distance (100 bp) to the nearest real TSS as TP, otherwise FP. And those real TSSs without a strong signal (>0.5) as a neighbor were counted as FN.

### Clustering and empirical p-value calculation

To reduce FDR by directly using outputs from the deep learning model, we developed a clustering-based method by grouping any sites with prediction score (outputted from the model and the range is from 0 to 1) larger than a pre-defined cutoff value C (default C = 0.1, we also tested 0.5 and 0.9, while 0.1 got the best performance) and within a pre-defined distance D (default D = 10bp, we also tested 30bp and 50bp, while 10bp had the best performance) into a cluster. The final score of each cluster was calculated by summing up prediction scores at each site within this cluster. Suppose that we have a cluster *c* that includes *n* sites, the final score *S_c_* was calculated based on the formula below.

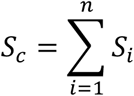

where *S_i_* is the prediction score outputted from the model in each site, and *S_i_* will be assigned to 0 if it is smaller than the defined *C*.

To further estimate an empirical p-value for each cluster, we made a null hypothesis that there was not any true TSS within the cluster, while the alternative hypothesis was that the cluster contained true TSS. The probability to reject the null hypothesis and accept alternative hypothesis is known as p-value, which was roughly calculated as shown below. The probability of a site *i* with score *S_i_* to be a true TSS was calculated by the total number of true TSSs with score above *S_i_* divided by the total number of TSSs with score above *S_i_* in the prediction. For instance, probability of a site with the prediction score 0.1 to be a true TSS, is around 3% among all sites with score above 0.1 from the three cell lines. For a cluster with *n* sites, the probability (*P_c_*) that it does not contain any true TSS was calculated below.

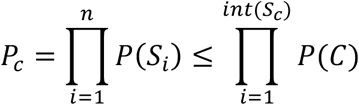

The *P*(*S_i_*) is the probability of a site with score *S_i_* that is not a true TSS and *int*(*S_c_*) is the round number of *S_c_*. Notably, a higher *S_i_* value will get a smaller *P*(*S_i_*), meanwhile *n* should be always no less than *int*(*S_c_*).

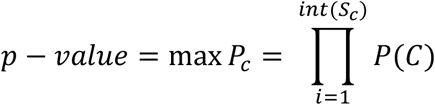

To simplify the calculation for each cluster, as well as to link the p-value to the final prediction score *S_c_*, we used the maximum probability as the p-value.

### Visualization of model features and identification of TFs

To understand sequence features learnt by our model during the fine-tuning stage of meta-learning, we paired each filter in the convolution layer of the meta-model and the fine-tuned model of each cell type, and calculated the motif gain of sequence features, defined as the difference of two positional weighted matrixes, among which each motif gain was compared with all transcription factor (TF) binding profiles of *Homo sapiens* in JASPAR (Fornes et al., 2020) to identify tissue-specific motifs and TFs. The matched motif-TF pairs with p-value < 0.001 were selected.

## Acknowledgements

We apologize to all colleagues whose work could not be cited due to space constraints. We thank all past and present members in Structural and Functional Bioinformatics (SFB) Group for their constructive feedback on this project. We also thank Mohammed Saif for providing generous support on computational resources.

## Author contribution

Juexiao Zhou: Conceptualization, Data curation, Methodology, Software, Writing - original draft, Visualization. Bin Zhan: Conceptualization, Data curation, Methodology, Software, Writing - original draft, Visualization. Haoyang Li: Data curation, Writing - review & editing. Longxi Zhou: Data curation, Writing - review & editing. Zhongxiao Li: Investigation, Writing - review & editing. Yongkang Long: Investigation, Writing - review & editing. Wenkai Han: Investigation, Writing - review & editing. Mengran Wang: Investigation, Writing - review & editing. Huanhuan Cui: Investigation, Writing - review & editing. Wei Chen: Investigation, Writing - review & editing, Supervision, Project administration, Funding acquisition. Xin Gao: Investigation, Writing - review & editing, Supervision, Project administration, Funding acquisition. All authors read and approved the final manuscript.

## Supplementary figure legends

**Supplementary Fig. S1:**
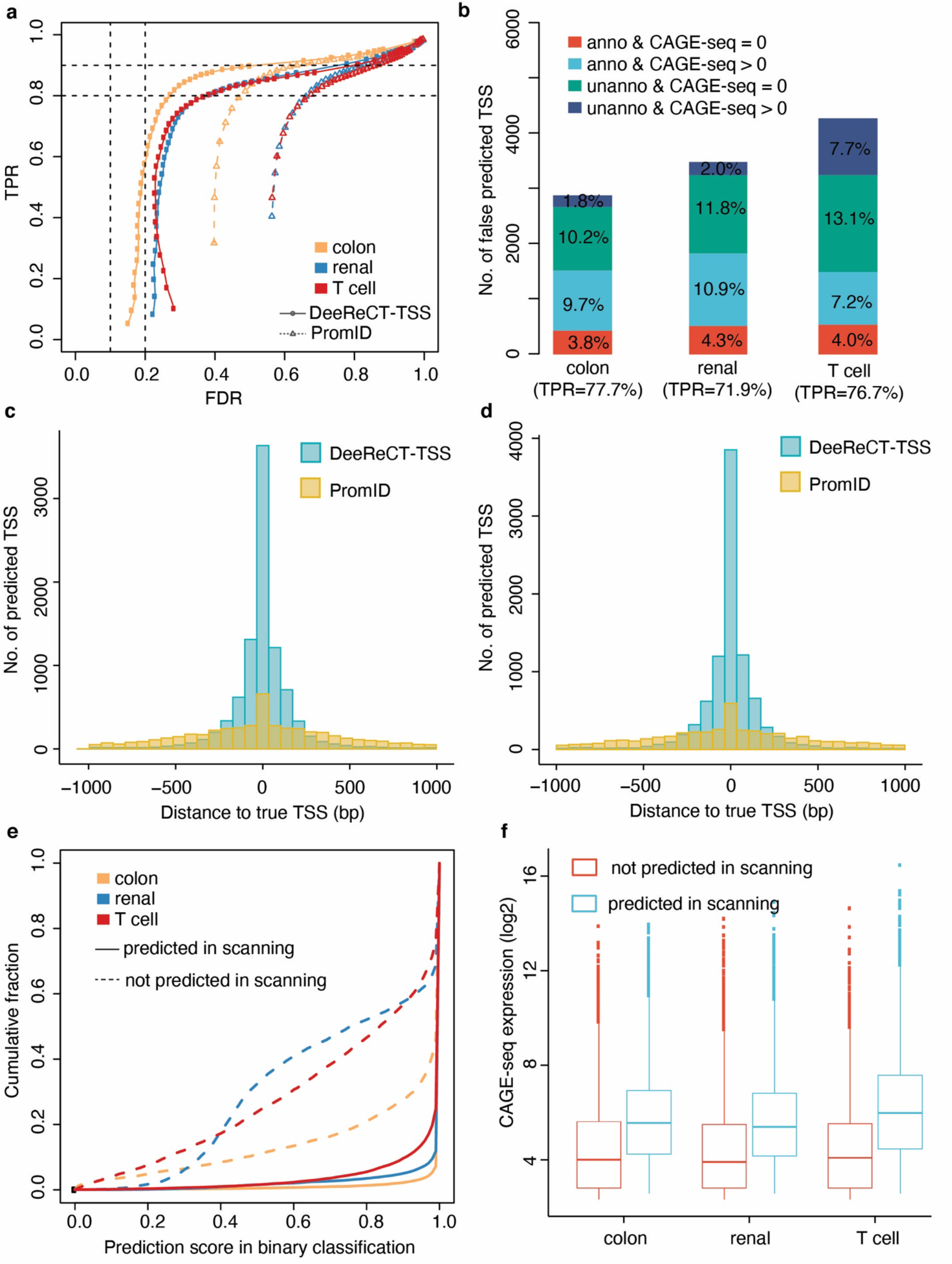
DeeReCT-TSS outperforms a recently published method in TSS identification. (a) Performance of DeeReCT-TSS and PromID in genome scanning after clustering the prediction score. (b) Barplot showing the number of four groups of false predicted TSS, including: 1, anno & CAGE-seq > 0, which is the TSS annotated in FANTOM and supported by CAGE-seq, but the expression did not pass the threshold; 2, anno & CAGE-seq = 0, which is the TSS annotated in FANTOM but not supported by CAGE-seq, 3, unanno & CAGE-seq > 0, which is the TSS not annotated in FANTOM, but supported by CAGE-seq, 4, unanno & CAGE-seq = 0, which is the TSS not annotated in FANTOM and not supported by CAGE-seq. Their percentages among the total predicted TSSs in each cell line were labeled in the bar. (c) and (d), Histogram of distance between predicted TSSs and true TSSs in renal (c) and T cell (d). (e) Cumulative plot showing the distribution of prediction score in binary classification for TSSs that are still successfully predicted in genome scanning or not in the three cell lines. (f) Expression level of TSSs that are successfully predicted in genome scanning and those are not in the three cell lines.

**Supplementary Fig. S2:**
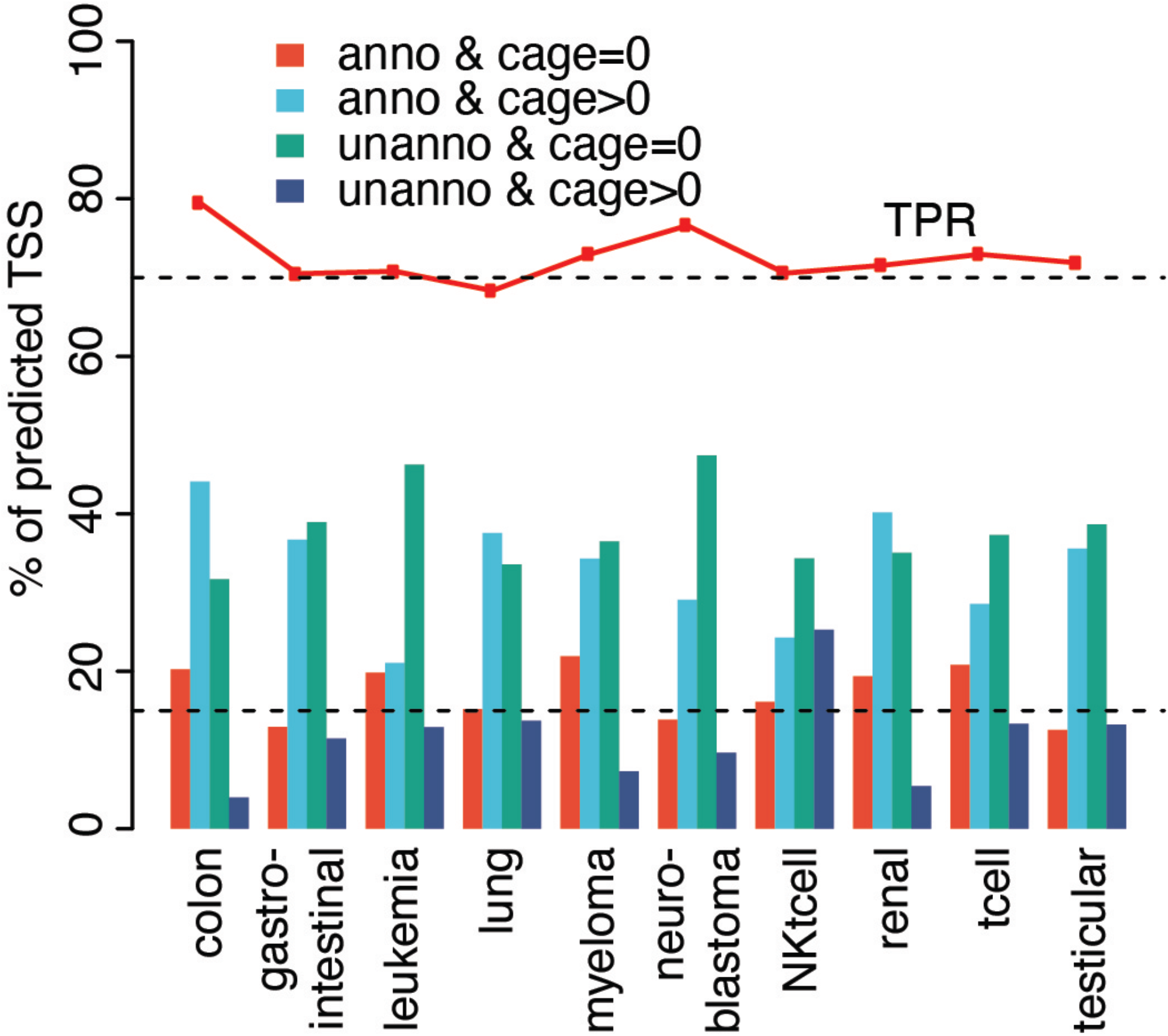
DeeReCT-TSS is capable of predicting cell-type-specific TSSs across 10 cell types. Barplot showing the percentage of four groups of false predicted TSSs. Four groups are the same as Supplementary Fig. S1b. The line above the bar indicates the TPR of the prediction in each cell line, while the dashed line indicates 70%.

**Supplementary Fig. S3:**
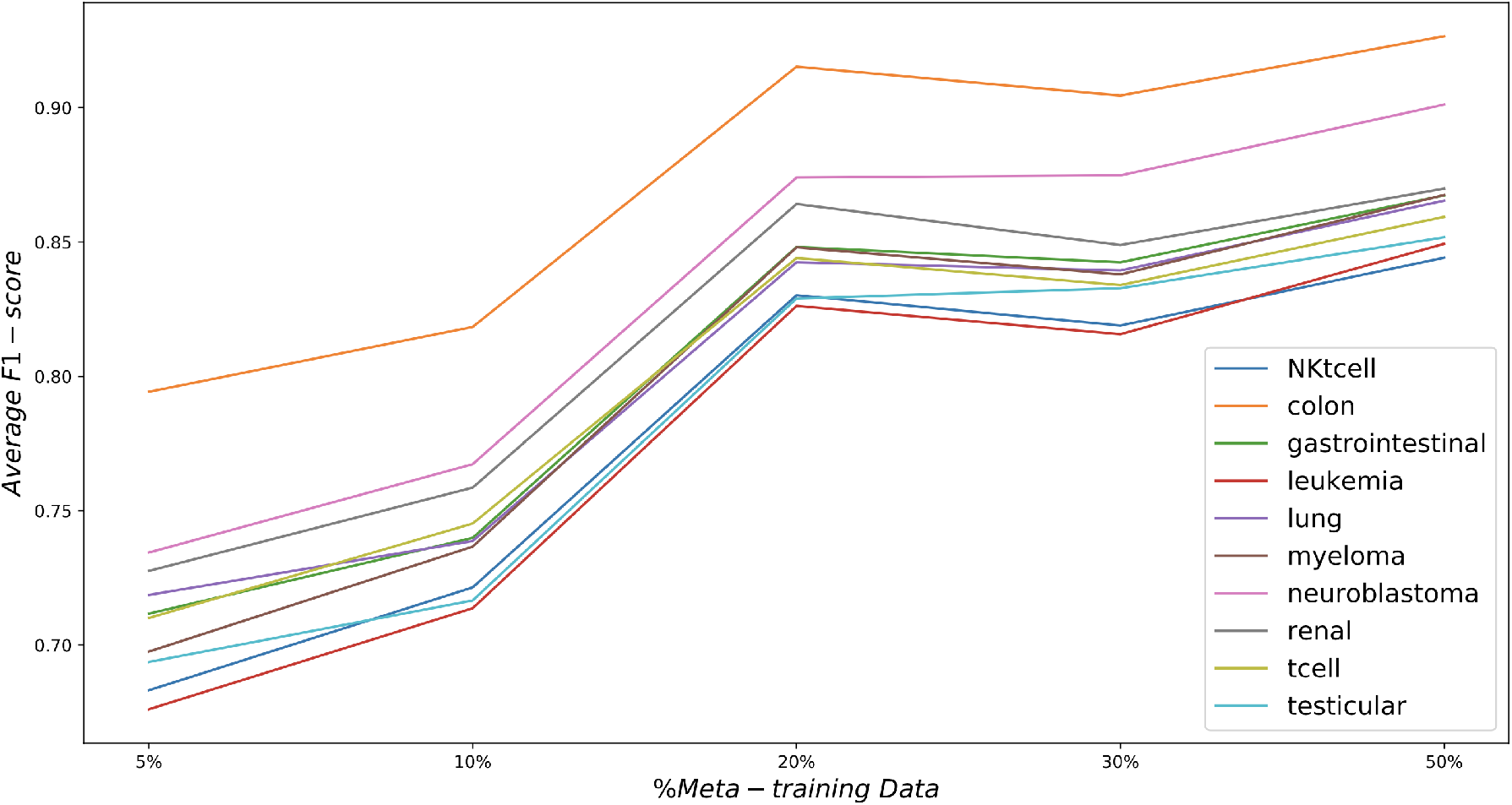
Ablation study showing performance of the fine-tuned model in 10 cell lines from meta-model trained using 5%, 10%, 20%, 30%, and 50% of TSSs in each cell line. X-axis is the proportion of TSSs used for meta-model in each cell line, y-axis is the average F1-score of the fine-tuned model.

